# Artificial selection of microbial communities to enhance degradation of recalcitrant polymers

**DOI:** 10.1101/474742

**Authors:** Robyn J. Wright, Matthew I. Gibson, Joseph A. Christie-Oleza

## Abstract

Recalcitrant polymers are widely distributed in the environment. This includes natural polymers, such as chitin, but synthetic polymers are becoming increasingly abundant, for which biodegradation is uncertain. Distribution of labour in microbial communities commonly evolves in nature, particularly for arduous processes, suggesting a community may be better at degrading recalcitrant compounds than individual microorganisms. Artificial selection of microbial communities with better degradation potential has seduced scientists for over a decade, but the method has not been systematically optimised nor applied to polymer degradation. Using chitin as a case study, we successfully selected for microbial communities with enhanced chitinase activities but found that continuous optimisation of incubation times between selective generations was of utmost importance. The analysis of the community composition over the entire selection process revealed fundamental aspects in microbial ecology: when incubation times between generations were optimal, the system was dominated by *Gammaproteobacteria*, main bearers of chitinase enzymes and drivers of chitin degradation, before being succeeded by cheating, cross-feeding and grazing organisms.

**Importance:** Artificial selection is a powerful and atractive technique that can enhance the biodegradation of a recalcitrant polymer and other pollutants by microbial communities. We show, for the first time, that the success of artificially selecting microbial communities requires an optimisation of the incubation times between generations when implementing this method. Hence, communities need to be transferred at the peak of the desired activity in order to avoid community drift and replacement of the efficient biodegrading community by cheaters, cross-feeders and grazers.

## 1 Introduction

Recalcitrant compounds are widely distributed in the environment (1–6). These include natural polymers, such as cellulose, (7) and chitin (1), and, more recently, xenobiotic compounds like plastics (2, 3, 5, 8), pesticides and detergents (9). Whilst processes to degrade natural compounds have had time to evolve and adapt, these processes may still require the participation of a consortia of organisms, each specialised in one of the multiple steps involved in the breakdown of the compound (10, 11). Laborious biodegradation processes are therefore rarely carried out entirely by a single microorganism in nature, and it is now well documented that a distribution of labour is favoured in natural microbial communities (12–16).

Many xenobiotic compounds have only existed in the last 50-100 years and microbial communities have had little time to evolve efficient biodegradation pathways to catabolise them. Some novel enzymes have, however, been discovered for new xenobiotic compounds, such as the recent discovery of an esterase involved in the degradation of poly(ethylene terephthalate) (PET) (17). This enzyme, termed “PETase”, is thought to have evolved from other esterases i.e. lipases and cutinases. Hence, although this enzyme shares considerable sequence homology with other enzymes capable of PET degradation (17–19), it has developed a higher hydrolytic activity against this polymer than any other tested esterase but, still, there is room for evolutionary improvement (18). The bacterium encoding this enzyme was isolated from a PET-degrading consortia of microorganisms and is capable of metabolising PET to its monomers, terephthalic acid and ethylene glycol (17). Similarly to the generation of toxic phenolic intermediates during lignin degradation (20), terephthalic acid can become toxic at high concentrations (21, 22) which suggests that degradation could be more efficient if carried out by a consortium rather than an individual organism. There are a number of examples of microbial consortia assembled to degrade recalcitrant xenobiotic compounds e.g. phthalic acid esters, benzene, xylene and toluene (23); polychlorinated biphenyls (24); polystyrene (25); and polyethylene (26) but, due to adverse biotic and abiotic conditions (e.g. temperature, humidity, competition and predation), the natural evolutionary development of novel biodegrading pathways and/or microbial consortia may be hampered in the environment (27).

Faster evolution can be achieved through artificial selection. A whole microbial community may be used as a unit of selection (artificial ecosystem selection) so that it becomes progressively better at a selective process over successive generations (28–31). The artificial selection of a measurable and desired trait is thought to outperform traditional enrichment experiments as it bypasses community bottlenecks and reduces stochasticity (31). Artificial ecosystem selection has been implemented for developing a microbial community capable of degrading the toxic environmental contaminant 3- chloroaniline (29) as well as to lower carbon dioxide emissions during growth (31) and generating a microbial community adapted to low or high soil pH (28). However, to our knowledge, it has not been previously used for improving polymer degradation and nor have the growth parameters involved (*e.g*. incubation time) been systematically optimised to enhance the selectivity of a desired process.

In the present study we aimed to optimise the artificial selection process of a marine microbial community for polymer degradation, using chitin as a case study. Chitin is one of the most abundant polymers on Earth (*i.e*. the most abundant polymer in marine ecosystems) constituting a key component in oceanic carbon and nitrogen cycles (1). Many microorganisms are already known to degrade chitin, and the enzymes and pathways used to do so are well characterised (10). We found that a microbial community could be artificially evolved to achieve higher chitinase activities, but there were certain methodological caveats to this selection process. We found that the incubation time between generations needed to be continuously optimised in order to avoid community drift and decay. Microbial community composition was evaluated and we confirmed that, if generation times are not continuously optimised, efficient biodegrading communities are rapidly taken over by cross feeders and predators with a subsequent loss of degrading activity.

## Results

### First artificial selection experiment; process optimisation

Our first artificial selection experiment highlighted the need to sub-culture each generation when the desired trait/chitinase activity was at its peak and not at a pre-defined incubation time, as done previously (28, 29, 31). Initially, we set a standardised nine-day incubation time for each generation because this was the time it took for chitinase activity to peak in a preliminary enrichment experiment (data not shown). After 14 generations we did not observe a strong increase in chitinase activity (Fig. 2A, and Supplementary Fig. S1) and, intriguingly, in nine out of the 14 generations we observed a lower activity in the positive selection than in the randomly selected control (Fig. 2A), suggesting that a random selection of microcosms is more effective in enhancing chitinase activity than actively selecting for the best communities. To further investigate the reasons behind this low efficiency, we took regular enzymatic activity measurements within generation 15 (Fig. 2B). As suspected, the chitinase activity was peaking much earlier within the generation, *i.e*. at day four, and by the end of the nine days the chitinase activity had dropped below the activities registered for the random selection experiment (Fig. 2B). Attending to this result, at generation 16 we set up an additional experiment, run in parallel, where the incubation time per generation was shortened to four days. Shortening the incubation time led to a selection of higher chitinase activities during generations 16 and 17, but the progressive increase in activity stalled by generations 18 and 19 (Fig. 2A). Chitinase activity was again measured daily within the final generation 20 and we found that the enzymatic activity was almost nine times higher on day two than day four (Fig. 2C), indicating that the optimal incubation time had again been reduced. While the nine-day incubation experiment gave an overall negative trend, shortening the incubation times to the chitinase maxima drastically increased the benefits of artificial community selection (Fig. 2A).

**Figure 1.**
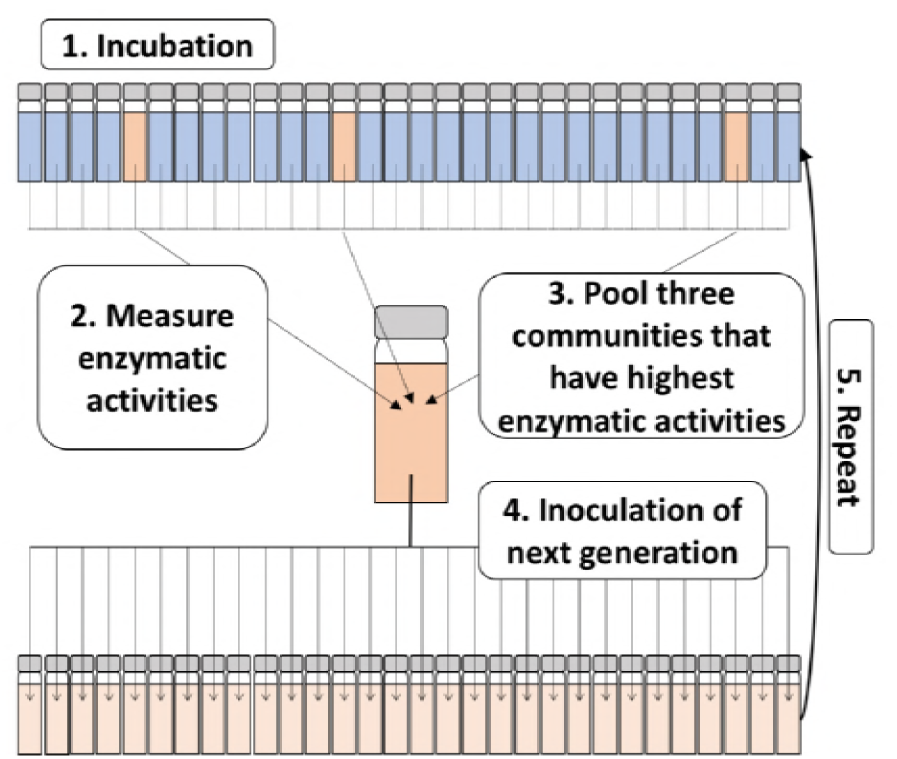
Method used for artificial selection of microbial communities. Briefly, 30 microcosms are inoculated with a natural community found in seawater (1). At the end of the incubation period, the enzymatic activity for a desired trait (*e.g*. chitinase activity) is measured for each microcosm (2). The three microcosms with the highest enzymatic activities are selected and pooled (3), and used to inoculate the next generation (4). This process is repeated over *n* generations (5).

**Figure 2.**
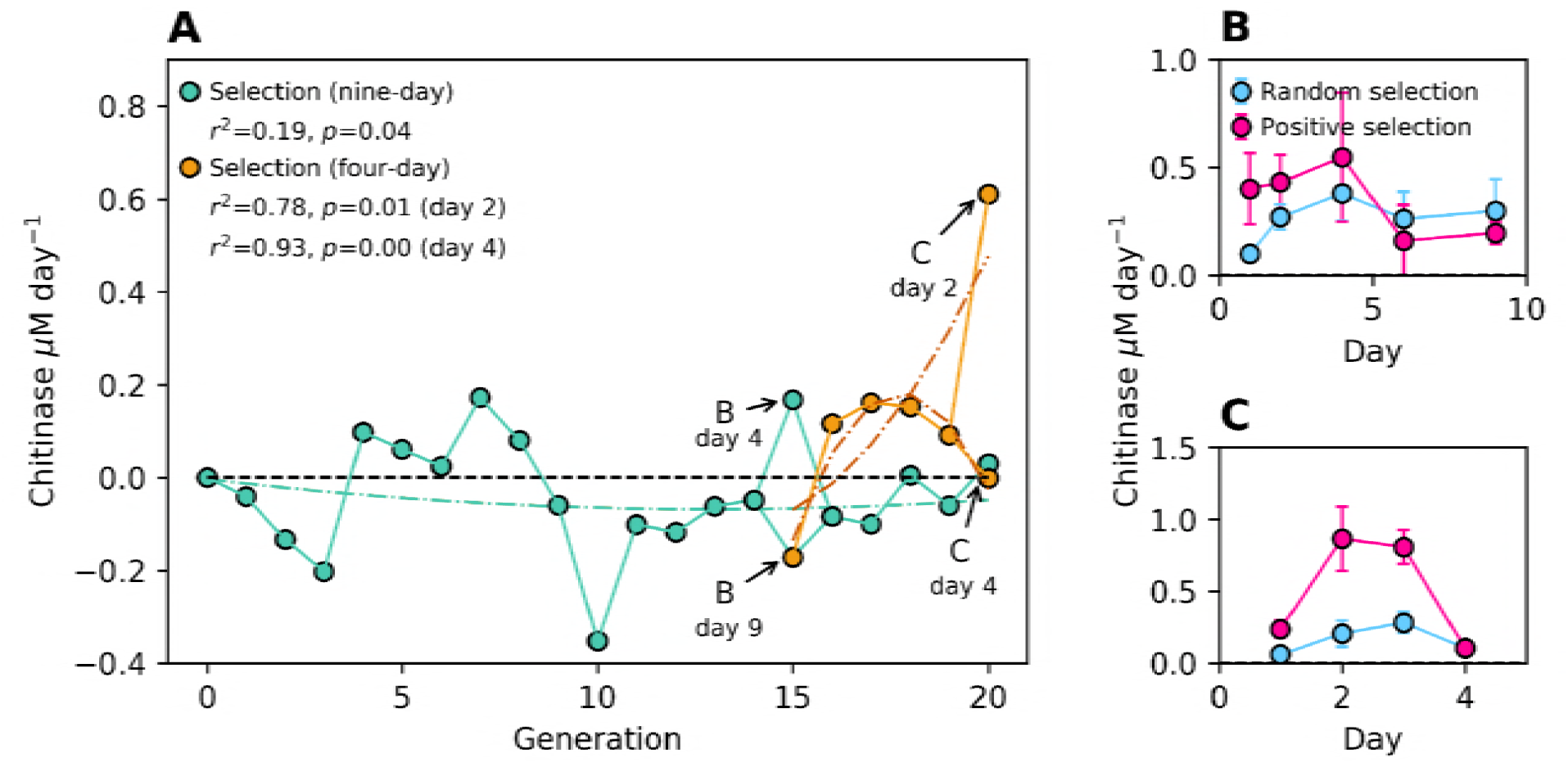
Chitinase activity in artificial selection experiment 1. (**A**) Enzymatic activity measured over 20 generations. Each point represents the mean of the positive selection communities (*n*=30) to which the mean of the randomly selected controls (*n*=30) was subtracted. The black dotted line (zero) represents where chitinase activity of the positive selection is equal to that of the random selection. Coloured dotted lines show the general trend for the respective incubation times generated by a linear regression model within Pythons’ statsmodels package (green and orange for nine- and four-day incubations, respectively). The *r^2^* and *p*-values for Pearson’s correlation coefficients between generation number and normalised chitinase activity are shown. (**B**) Chitinase activity measured within generation 15 of the 9-day incubation. (**C**) Chitinase activity measured within generation 20 of the 4-day incubation. In panels B and C, each point represents absolute chitinase activity measured in the positive (red) and random selection (blue).

### Microbial community succession

We carried out MiSeq amplicon sequencing of the 16S and 18S rRNA genes to characterise the microbial community succession that occurred within the first selection experiment and, by this way, gain insight into the strong variability in chitinase activity observed over time. We sequenced the communities that were used as the inoculum of each of the 20 generations, both nine and four-day long experiments, as well as the community obtained from the daily monitoring of generation 20. This data was processed using both Mothur (32) and DADA2 workflows (33, 34), obtaining similar results (Supplementary Figs. S2 and S3). DADA2 results are presented here as this workflow retains greater sequence information, better identifies sequencing errors and gives higher taxonomic resolution (35). Unique taxa are therefore called amplicon sequence variants (ASVs) rather than operational taxonomic units (OTUs).

#### Community succession over the four-day incubation period within generation 20

The daily microbial community analysis over the four days at generation 20 showed a progressive increase in prokaryotic diversity (from 0.83 to 0.93, according to Simpsons index of diversity) whereas a strong decrease in diversity was observed amongst the eukaryotic community (from 0.93 to 0.38; Fig. 3A). SIMPER analyses were carried out to identify those 16S and 18S rRNA gene ASVs that were contributing most to the differences over the four successive days observed in Fig. 3B. The top five ASVs in these analyses were responsible for 50% and 60% of variation for the 16S and 18S rRNA genes, respectively (Fig. 3C).

**Figure 3.**
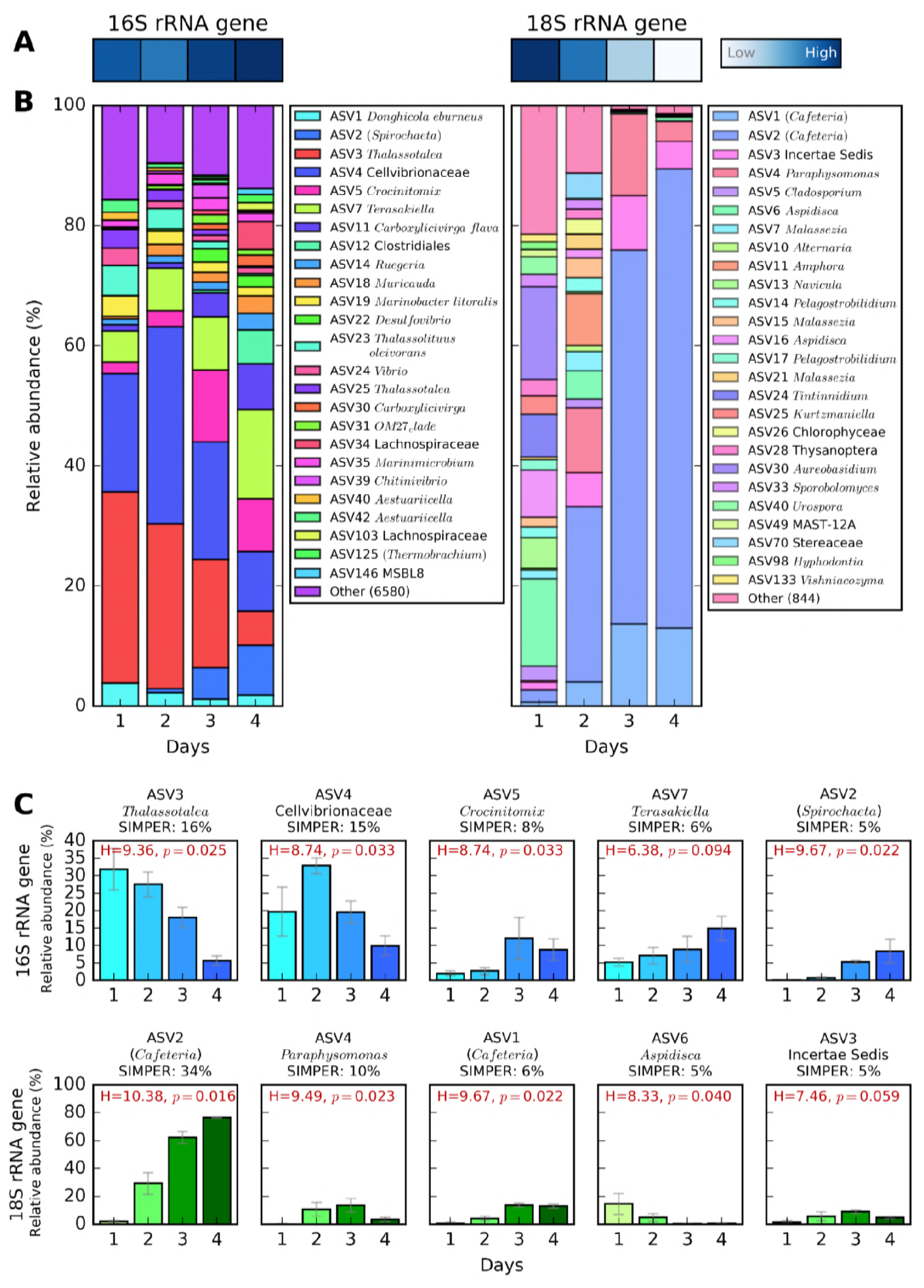
Daily microbial community analysis over the four day incubation period within generation 20. The analysis was performed on the three communities that showed highest chitinase activity by the end of the four days and which would have been used to inoculate the next generation. (**A**) Simpsons index of diversity of the 16S (left) and 18S rRNA gene (right) amplicon analysis. Scale ranges between 0.38 (low) and 0.93 (high). (**B**) Community relative abundance over the 4 day incubation period. Only ASVs with abundance above 1% in at least one time point are shown. The abundance for each ASV is a mean value from the three communities. ASVs were classified to genus level by SILVA. Names in brackets were not identifiable with the standard analysis pipeline and were identified through a BLAST search of the NCBI database. (**C**) Five 16S and 18S rRNA gene ASVs that contributed the most to the community variations over time according to a SIMPER analysis. The percentage of variation to which each ASV contributes is indicated. Error bars represent the standard deviations of three communities used to inoculate the next generation.

For the 16S rRNA gene, the most important ASVs were: ASV3 (*Thalassotalea*, contributing to 16% of the community variation between the four days, *p*=0.025), ASV4 (Cellvibrionaceae, 15% variation, *p*=0.033), ASV5 (*Crocinitomix*, 8% variation, *p*=0.033), ASV7 (*Terasakiella*, 6% variation, *p*=0.094) and ASV2 (*Spirochaeta*, 5% variation, *p*=0.022) (Fig. 3C). ASVs 3 and 4 (both *Gammaproteobacteria*) represented over 50% of the prokaryotic community abundance on day 2, when chitinase activity was highest, and their abundances followed a similar pattern to the chitinase activity over the four days (Fig. 2C), suggesting that these ASVs may be the main drivers of chitin hydrolysis. On the other hand, ASVs 7 and 2 both showed a progressive increase over time (*i.e*. from a combined relative abundance of 5% on day 1 to 23% on day 4; Fig. 3C), suggesting that these ASVs could be cross- feeding organisms that benefit from the primary degradation of chitin. Interestingly, the overall 16S rRNA gene analysis also showed a strong succession over time at higher taxonomic levels (Fig. 4). While *Gammaproteobacteria* pioneered and dominated the initial colonisation and growth, presumably, via the degradation of chitin (*i.e*. with 73% relative abundance during the first two days), all other taxonomic groups became more abundant towards the end of the incubation period (*e.g. Clostridia*, *Bacteroidia* and *Alphaproteobacteria* increased from an initial relative abundance of 0.1, 2.8 and 12% on day one to 13.5, 22 and 21% on day four, respectively; Fig. 4). Microbial isolates confirmed *Gammaproteobacteria* as the main contributors of chitin-biodegradation (as discussed below).

**Figure 4.**
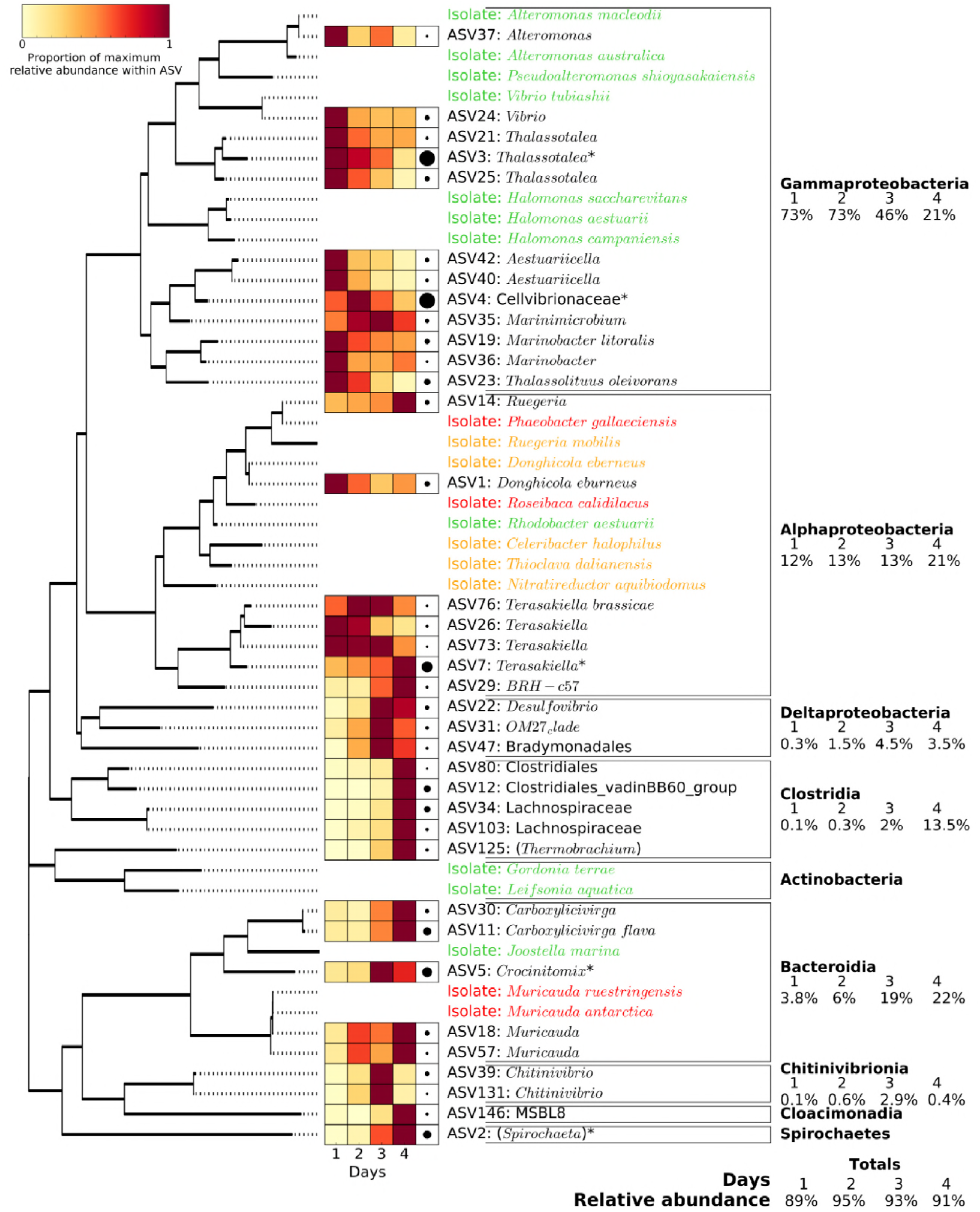
Phylogenetic analysis and relative abundance of the major 16S rRNA gene ASVs (*i.e*. with relative abundance above 0.5% in at least one of the four days) and bacterial isolates obtained at the end of the artificial selection experiment. Phylogenetic grouping is represented by a mid-point rooted maximum likelihood phylogenetic tree. The 36 ASVs represented in the figure (out of the 6605 total ASVs detected) accounted for 92% of all 16S rRNA gene relative abundance. The heatmap represents the relative abundance of each ASV over the four days, with darker red showing the day at which the ASV showed maximum abundance. Black circles on the right of the heatmap represent the maximum relative abundance for that ASV amongst the entire community. The 20 isolates are coloured depending on their ability to grow on chitin and the monomer, GlcNAc (green), the GlcNAc only (orange), or neither (red).

The SIMPER analysis of the 18S rRNA gene highlighted ASV2 (*Cafeteria* sp., contributing to 34% of the community variation between the four days, p=0.016), ASV4 (*Paraphysomonas*, 10% variation, p=0.023), ASV1 (*Cafeteria* sp., 6% variation, p=0.392), ASV6 (*Apsidica*, 5% variation, p=0.040) and ASV3 (Incertae Sedis, 5% variation, p=0.059) as the five main ASVs contributing to 60% of the community variation over the four days (Fig 3C). ASV2, which was 96% similar to the bactivorous marine flagellate *Cafeteria* sp., was by far the most striking Eukaryotic organism, showing an increase in relative abundance from 2% on day 1 up to over 76% on day 4 (Fig. 3B and 3C). As observed in prokaryotes, Eukaryotic phylogenetic groups also showed a large variation between the beginning and the end of the incubation period, mainly due to the increase of *Bicosoecophyceae* over time (*i.e*. from 2.6 to 89% relative abundance driven by both ASV1 and ASV2; Supplementary Fig. S4).

#### Community succession over the entire artificial selection experiment

We analysed the 16S and 18S rRNA gene community composition (Supplementary Fig. S5) at the end of each generation in order to determine the effect that positive or random selection of communities had across the 20 generations, both for the nine-day incubation experiment (*i.e*. generations 0 to 20) and shortened four-day incubation experiment (*i.e*. generations 16 to 20). Most interestingly, the overall community variability across all generations (16S and 18S rRNA gene nMDS analysis; Fig. 5A) showed that only the positive selection of the shortened four-day incubations differentiated the community from the random selection, which was confirmed by a PerMANOVA test using Bray-Curtis distance (16S rRNA gene *p*=0.001; 18S rRNA gene *p*=0.002; Supplementary Table S2), while the nine- day selection mostly clustered with the random control communities. This is a clear explanation as to why the nine-day incubation time was not allowing a progressive selection of a community with better chitinase activities than those obtained randomly and, only when the time was shortened, did we observe an effect of the positive selection over the random selection.

**Figure 5.**
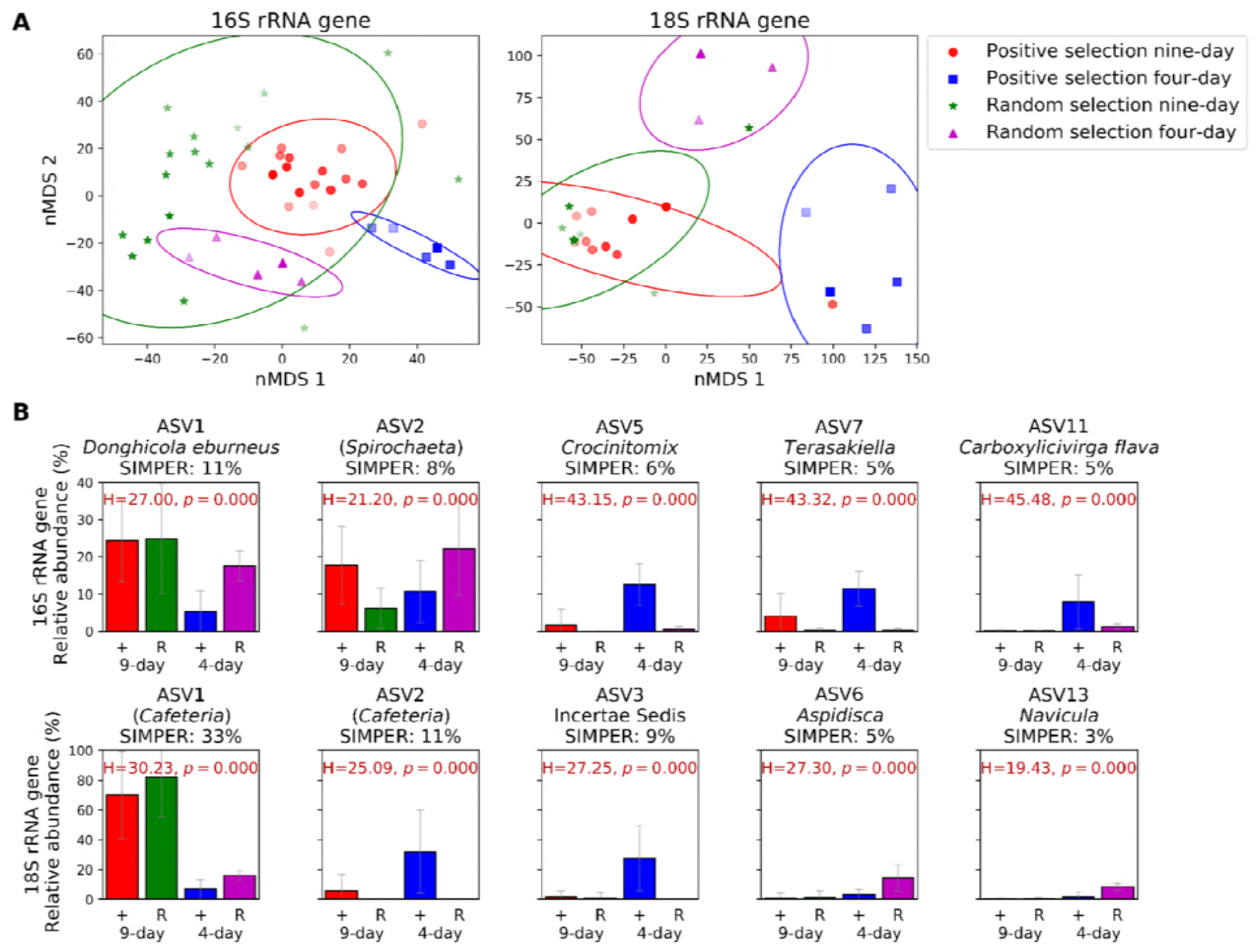
Microbial community variation over the entre artificial selection experiment. (**A**) nMDS plot showing Bray-Curtis distance of 16S (left) and 18S communities (right). Distance between the community composition obtained from nine-day (red circles) and four-day incubations (blue squares) of the positive selection, and nine-day (green stars) and four-day incubations (purple triangles) of the random controls are shown. Marker colour intensity correlates to generation number, where progressive darker colours represent later generations. Each point represents the mean of the three communities selected from one generation used to inoculate the following one. Ellipses show the mean plus the standard deviation of each group of samples. Stress values are 0.175 for the 16S rRNA gene and 0.063 for the 18S rRNA gene. (**B**) Five 16S (top panel) and 18S rRNA gene ASVs (bottom panel) that contributed the most towards community variations between the nine-day (generations 0-20) and four-day (generations 16-20) positive (+) and random (R) selections according to SIMPER analyses. The percentage of variation to which each ASV contributes is indicated. ASVs were classified to the species level with the standard analysis pipeline using the SILVA database where possible. Names in brackets were not identifiable and were identified through a BLAST search of the NCBI database. Relative abundances and error bars shown are the mean and standard deviations of all generations within that treatment.

SIMPER analyses were carried out to determine the ASVs that most strongly contributed to the differences between groups (*i.e*. positive *versus* random selections and nine-day *versus* four-day incubation times; Fig. 5B). For the 16S rRNA gene, the top five ASVs identified by the SIMPER analysis contributed to 35% of the community variation, while for the 18S rRNA gene, they accounted for 61% (Fig. 5B). The 16S rRNA gene ASVs 5, 7 and 11 (*Crocinitomix*, *Terasakiella* and *Carboxylicivirga flava*, respectively) presented a much higher abundance in the four-day positive selection than in any other selection (13%, 11% and 8%, respectively), suggesting that these species were the major contributors to the differentiation of these communities, as seen in Fig. 5A. As observed above for the four-day incubation analysis, *Cafeteria* sp. (18S rRNA gene ASV1 and ASV2, both 96% similar) was again the most conspicuous Eukaryotic organism. ASV2 was more abundant in the positive four-day selection (32% of the relative abundance), while ASV1 was highest in the three other selections (70% and 82% in the positive and random nine-day selection, respectively, and 16% in the random four-day selection; Fig. 5B).

#### Chitinase gene copies in artificially-assembled metagenomes

Artificially-assembled metagenomes, generated by PICRUSt (36) from the 16S rRNA gene amplicon sequences, were used to search for enzymes involved in chitin degradation: KEGG orthologs K01183 for chitinase, K01207 and K12373 for chitobiosidase, K01452 for chitin deacetylase, and K00884, K01443, K18676 and K02564 for the conversion of GlcNAc to Fructose-6 phosphate (Supplementary Fig. S6) (37–39). As expected from the measured chitinase activities, the shortened four-day incubation experiment showed over 30 times more chitinase (K01183) gene copies than the nine-day incubation experiment (*i.e*. an average of 0.66 copies per bacterium were observed in the four-day incubation experiment while only 0.025 copies per bacterium were observed over the same generations in the nine-day experiment). Also, from the daily analysis of generation 20, the chitinase activity was positively correlated with the normalised chitinase gene copy number (*r2*=0.57), with a peak in chitinase activity *and* chitinase gene copies on day 2 (*i.e*. over one chitinase gene copy per bacterium). The most striking result from this analysis was the strong bias of taxonomic groups that contributed to the chitinase and chitin deacetylase genes; chitinase genes were mainly detected in *Gammaproteobacteria* and some *Bacteroidia*, whereas the chitin deacetylase genes were almost exclusively present in *Alphaproteobacteria*. It is worth highlighting that the chitosanase gene (K01233), the enzyme required to hydrolyse the product from chitin deacetylation, chitosan, was not detected in any of the artificial metagenomes. Chitobiosidases (K01207 and K12373) and enzymes involved in the conversion of GlcNAc to Fructose-6 Phosphate (K00884, K01443, K18676 and K02564) were more widespread. Nevertheless, this data needs to be taken with caution as these were not real metagenomes.

#### Isolation and identification of chitin degraders

Bacterial isolates were obtained from the end of the artificial selection experiments to confirm the ability of the identified groups to degrade chitin. From the 50 isolates obtained, 20 were unique according to their 16S rRNA gene sequences. From these, 18 showed at least 98% similarity with one or more of the MiSeq ASVs (Supplementary Table S3) although, unfortunately, none belonged to the most abundant ASVs detected during the community analysis. The ability for chitin and GlcNAc degradation by each one of the isolates was assessed. We found that 16 of these isolates could grow using GlcNAc as the sole carbon source, but only 11 of these strains could grow on chitin (Fig. 4). The four remaining bacteria from the 20 isolated could not grow using chitin or GlcNAc. Most interestingly, all isolates from the class *Gammaproteobacteria* (*n*=7) were capable of chitin degradation whereas only a smaller subset of isolates had this phenotype in other abundant taxonomic groups, such as *Bacteroidia* (1 out of 3) or *Alphaproteobacteria* (1 out of 8; Fig. 4).

### Second artificial selection experiment; implementing an improved selection process

A second selection experiment showed an extremely rapid boot in chitinase activity *i.e*. reaching almost 90 µM day^−1^ in only 7 generations (Figs. 6, and S7), when the maximum activity achieved in the first experiment was 0.9 µM day^−1^ (Fig. 2C), demonstrating that implementing an optimised incubation time between generations largely enhances the selection of a desired trait. Chitinase activity was measured daily until a peak in chitinase activity was observed. The communities with highest chitinase activity on this day were used to start the next generation.

**Figure 6.**
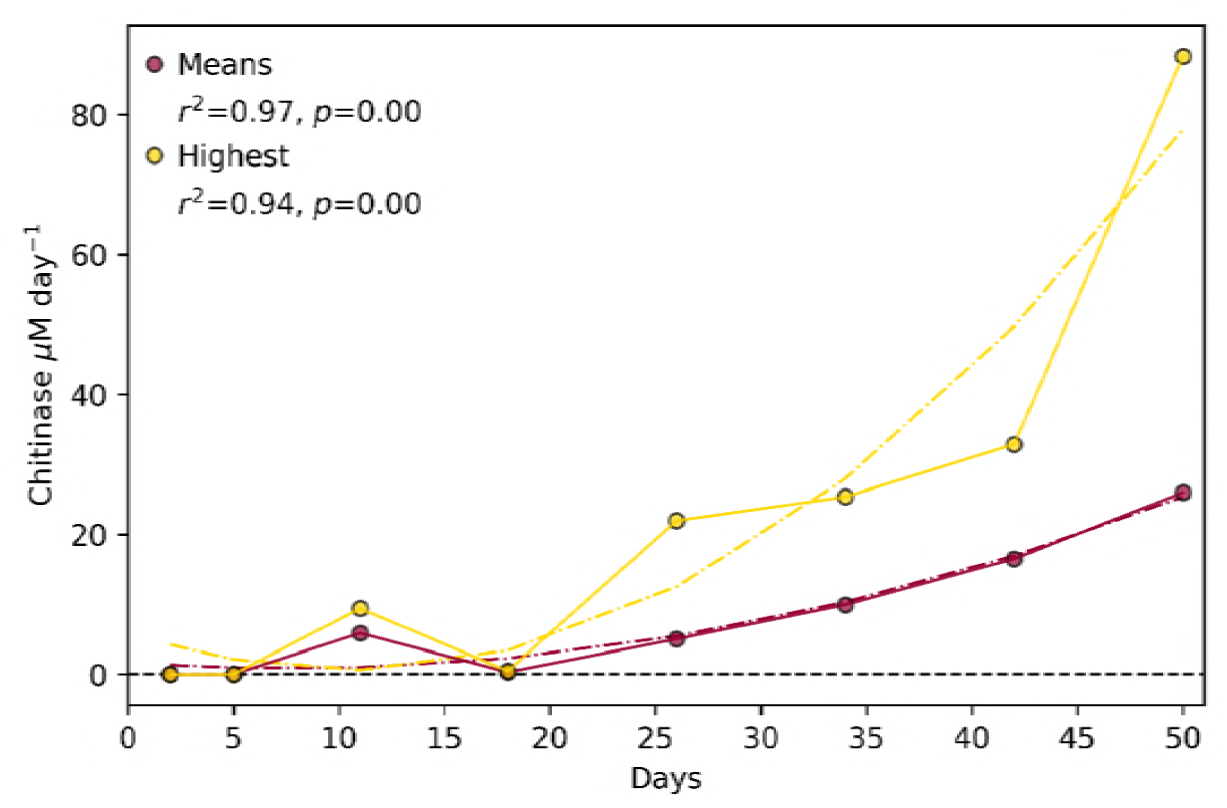
Chitinase activity of artificial selection experiment 2. Graph show the mean chitinase activity of the positive selection, from which the mean random selection was subtracted. The means of all communities within the generation (*n*=30; red) and those of only the three communities that were pooled for the inoculum of the next generation (yellow) are shown. The *r2* and *p*-values are for Pearson’s correlation coefficients and lines of best fit (dotted lines) were determined using linear regression models within Pythons’ statsmodels package.

## Discussion

Artificial selection of microbial communities is, in principle, a powerful and atractive technique which has surprisingly been used in only a limited number of studies to date (28, 29, 31), possibly due to the lack of success as a consequence of poor process optimisation. Here, using chitin degradation as a case study and a detailed analysis of the community succession, we show that artificial selection of microbial communities can be largely improved by controlling the incubation times between generations. The rapid succession of microbial community structure means generations need to be transferred at the peak of the selected phenotypic activity (*e.g*. chitinase activity) or these get rapidly replaced by less efficient communities of cross-feeding microorganisms (*i.e*. bacteria and grazers). Previous studies that have artificially selected microbial communities for a particular phenotype did not report optimisation of the incubation time between generations (28, 29, 31) which, in our hands, would have resulted in a negative selection (Fig. 2). In agreement with our results, Penn and Harvey (2004) (40) suggested that the observed phenotype in artificial ecosystem selection experiments could be significantly affected by community structure.

An understanding of microbial ecology helps explain the importance of the timing during generation transfer. Datta et al. (2016) (42) observed three distinct stages of community structure during the colonisation of chitin particles: (a) attachment; (b) selection, and; (c) succession. Each phase was characterised by having relatively higher abundances of organisms that were: (a) good at attaching to chitin particles; (b) good at degrading chitin particles, and; (c) not able to degrade chitin, but able to benefit from others that could, *i.e*. “cheaters” and cross-feeders (43, 44). During our first experiment, as communities become better and faster at degrading chitin, we were measuring the chitinase actvity when the communities were in the succession rather than in the selection stage and, therefore, the active chitinolytic community had decayed and was dominated by cheaters and cross-feeders (Figs. 3 and 4). Hence, it was only when selecting at phenotypic time optima when chitinase activity improved and the overall community differentiated from the random control communities (Fig. 5 and 6). It is also interesting to note the selection of the grazer *Cafeteria* sp. (90% of the Eukaryotic community), a genus of bactivorous marine flagellates that are commonly associated with marine detritus (45). The predator-prey dynamics postulated by Lotka–Volterra’s equations would also support the need to shorten generation times to favour the prey’s growth *i.e*. chitinolytic bacteria (46, 47).

Interestingly, a strong successional pattern was observed at a higher taxonomic level. While *Gammaproteobacteria* dominated during the initial stages when chitinase activity was at its peak (accounting for over 70% of the prokariotic community), other groups increased in abundance during the later stages (*i.e. Alphaproteobacteria*, *Bacteroidia* and *Clostridia*), similarly to the pattern previously observed by Datta et al. (2016) (42).

The fact *Gammaproteobacteria* are major contributors to chitin degradation is not new (48–53). All *Gammaproteobacteria* isolates obtained from the end of the experiments were able to grow using chitin as the only source of carbon and energy (Fig. 4) confirming that this class is likely responsible for most of the chitinase activity observed. On the other hand, *Alphaproteobacteria*, the numerically dominant class of heterotrophic bacteria in surface oceans (54, 55), follow a cross-feeding and/or cheating life-strategy as five out of eight *Alphaproteobacterial* isolates could only use N-acetyl-D-glucosamine (GlcNAc) and only one could use chitin. This was confirmed by the PICRUSt metagenome analysis (Fig. S6), where almost all chitinase enzymes copies were encoded by *Gammaproteobacteria* (*i.e*. 90%; encoding almost one gene copy per bacterium) and, to a lesser extent, by some *Bacteroidia*. Chitin is made up of molecules of GlcNAc linked by (1,4)-β-glycosidic bonds, and it has previously been found that initial degradation of chitin takes place predominantly by: i) chitinases which depolymerise the (1,4)-β-glycosidic bonds either at the ends or in the middle of chains, or ii) chitobiosidase enzymes which also hydrolyse (1,4)-β-glycosidic bonds but only at the ends of chitin chains. Genes for the intracellular enzymes involved in GlcNAc utilisation (*i.e*. transformation of GlcNAc to Fructose-6-phosphate) were much more widespread amongst different taxonomic groups, highlighting the broader distribution of cross-feeding or cheating organisms which can benefit from the extracellular depolymerisation of chitin which generates freely available GlcNAc to the community. Alternative degradation of chitin may also occur by deacetylation and deamination of the GlcNAc amino sugar, transforming chitin into chitosan and cellulose, respectively, after which they can be depolymerised by a range of other enzymes (*e.g*. chitosanases or cellulases) (10, 56, 57). While *Alphaproteobacteria* did not contribute to chitinase enzymes, it *did* potentially encode for most of the chitin deacetylases in the system, although no chitosanases were detected.

Chitinolytic organisms have previously been found to make up between 0.1 and almost 6% of prokaryotic organisms in aquatic ecosystems (43, 58), while over a third of the organisms in these habitats can utilise only the products of chitin hydrolysis (*i.e*. GlcNAc) (43, 59–61). With *Gammaproteobacteria* being primarily responsible for the degradation of chitin here, the success of the artificial selection for an enhanced chitinolytic community was possibly achieved by the selective enrichment of this group between the beginning (5% of the prokaryotic community, within the expected range of *Gammaproteobacteria* found within natural environments) (43, 58) and end of the experiment (75% of the community).

Here we have proven the validity of artificially selecting a natural microbial community to better degrade a recalcitrant polymer, but have highlighted the caveats for achieving this goal, which require a better understanding of the ecology of the system. We found that optimisation of incubation times is essential in order to successfully implement this process, as optimal communities enter rapid decay due to their replacement by cheaters and cross-feeders, as well as the increase of potential predators such as grazers and, although not tested here, viruses. Hence, future artificial selection experiments should adjust generation incubation times to activity maxima to successfully evolve enhanced community phenotypes.

## Materials and methods

### Microbial inoculum

The microbial community used as an inoculum was obtained from bulk marine debris collected during boat tows from both Plymouth Sound (Devon, UK; June 2016) and Portaferry (Northern Ireland, UK; August 2016).

### Chitinase activity measurements

Chitinase activity was measured as the liberation of the fluorogescent molecule 4-methylumbelliferyl (MUF) from three chitinase substrates (MUF-N-acetyl-β-D-glucosaminide, MUF-β-D-N,N’- diacetylchitobioside and MUF-β-D-N,N’,N’’-triacetylchitotrioside; Sigma Aldrich, UK), following the previously described method (49, 62, 63) (Supplementary information). Standards curves were obtained using chitinase from *Streptomyces griseus* (Sigma Aldrich, UK) dissolved in sterile phosphate buffered saline solution (pH 7.4; 0.137 M) with a highest concentration of 0.1 U mL^−1^ (activity equivalent to 144 µM day^−1^). Samples were diluted prior to measurement if they were expected to be above this range.

### Artificial selection

The process for artificial selection is depicted in Figure 1. Briefly, 30 individual microcosms per treatment and generation were incubated in the dark under the conditions described below. At the end of each generation the three microcosms with the highest chitinase activities (or three random microcosms in the case of the control) were pooled and used as the inoculum for the next generation of microcosms (*n*=30). This was repeated across multiple generations. Two artificial selection experiments were performed, the first to optimise the process, and the second to implement optimal conditions and achieve a high-performing chitinolytic microbial community:

#### First artificial selection experiment

Incubations were carried out at 23°C in 22 mL glass vials (Sigma Aldrich), each containing 20 mL of autoclaved seawater (collected from outside Plymouth Sound, Devon, UK; June 2016) supplemented with NaH_2_PO_4_, F/2 trace metals (64) (Supplementary information) and 100 mg of chitin powder (from shrimp shells; Sigma Aldrich) as the sole source of carbon and nitrogen. Generation 0 was started with 200 µL of microbial inoculum. The efficiency of the selection process was assessed by comparing a ‘positive selection’ (where the three communities with highest activity were pooled and 200 µL was used to inoculate each one of the 30 microcosms of the next generation) against a ‘random selection’ (where three communities were chosen at random, using a random number generator within the Python module Random, to inoculate the following generation) to give a control against uncontrollable environmental variation (65). Each treatment was repeated across 20 generations with incubation times of nine days. In parallel, treatments where incubation times were shortened to four days were setup after generation 15. Samples were taken from each community and stored in 20% glycerol at −80°C for further microbial isolation, and pellets from 1.5 ml of culture were collected by centrifugation (14,000 *x g* for 5 mins) and stored at −20°C for final DNA extraction and community analysis.

#### Second artificial selection experiment

A second selection experiment was setup implementing optimal generation times. Microcosms were incubated in 2 mL 96-well plates (ABgene , ThermoFisher Scientific) covered by Corning^®^ Breathable Sealing Tapes to stop evaporation and contamination while allowing gas exchange. Each well contained 1.9 mL of a custom mineral media containing MgSO_4_, CaCl_2_, KH_2_PO_4_, K_2_HPO_4_, 0.52 M NaCl and artificial seawater trace metals (Supplementary information), supplemented with 10 mg of chitin powder. The microbial inoculum was 100 µL (*i.e*. initial inoculum and transfer between generations). Chitinase activity was measured daily. Transfer between generations was carried out just after the peak of chitinase activity had occurred, calculated as the average chitinase activity across the 30 microcosms of the positive selection treatment. Plates were incubated in the dark at 30°C with constant shaking (150 rpm). Eight days was the maximum incubation time allowed to reach maximum chitinase activity due to volume constraints.

### DNA extraction and amplicon sequencing

DNA was extracted using the DNeasy Plant Mini Kit (Qiagen) protocol, with modifications as follows (adapted from 66): 300 µL 1 × TAE buffer was used to resuspend cell pellets and these were added to ~0.4 g of sterile 0.1 mm Biospec zirconia silica beads in 2 mL screw cap microtubes (VWR international). Bead beating was carried out for 2 × 45 s and 1 × 30 s at 30 Hz using a Qiagen Tissue Lyser. Cell lysates were then processed in accordance with the manufacturer’s instructions, with an extra centrifugation step to ensure all liquid was removed (1 min, 13,000 *× g*) directly before elution of samples. A Qubit^®^ HS DNA kit (Life Technologies Corporation) was used for DNA quantification after which they were diluted to equalise the concentrations across samples. A Q5^®^ Hot Start High-Fidelity 2X Master Mix (New England Biolabs^®^ inc.) was used to amplify the 16S rRNA gene v4-5 regions using primers 515F-Y and 926R (67), and the 18S rRNA gene v8-9 regions using primers V8F and 1510R (68) (Supplementary information). PCR products were purified using Ampliclean magnetic beads (Nimagen, The Netherlands). Index PCR was carried out using Illumina Nextera Index Kit v2 adapters. Samples were normalised using a SequelPrep™ Normalisation Plate Kit (ThermoFisher Scientific). Samples were pooled and 2 × 300 bp paired-end sequencing was carried out using the MiSeq system with v3 reagent kit. Negative and chitin only DNA extraction controls and library preparation negative controls were processed and sequenced alongside samples.

### Microbial community structure determination

Two different workflows were used to analyse the sequencing data: DADA2 (33, 34) and Mothur (32). DADA2 delivers better taxonomic resolution than other methods (*e.g*. Mothur) as it retains unique sequences and calculates sequencing error rates rather than clustering to 97% similarity (35). The resultant taxonomic units are referred to as amplicon sequence variants (ASVs) rather than operational taxonomic units (OTUs from Mothur). For the DADA2 analysis, sequencing data were processed following the DADA2 (version 1.8.0) pipeline (33). Briefly, the data were filtered, *i.e*. adapter, barcode and primer clipped, and the ends of sequences with high numbers of errors were trimmed. The amplicons were denoised based on a model of the sequencing errors and paired end sequences were merged. Only sequences between 368 – 379 for the 16S rRNA gene and 300 – 340 for the 18S rRNA gene were kept and chimeras were removed. The resulting ASVs were classified using the SILVA reference database (v132) (69). For the Mothur analysis (32), sequencing data were filtered *i.e*. adapter, barcode and primer clipped, sequence length permitted was 450 bp for the 16S rRNA gene and 400 bp for the 18S rRNA gene, maximum number of ambiguous bases per sequence = 4, maximum number of homopolymers per sequence = 8. Taxonomy assignment was performed using the SILVA reference database (Wang classification, v128) (69) and operational taxonomic units (OTUs) set at 97% similarity. For both processing workflows, chloroplasts, mitochondria and Mammalia were removed from the 16S rRNA gene and 18S rRNA gene datasets, eukaryotes were removed from the 16S rRNA gene dataset, and bacteria and archaea from the 18S rRNA gene dataset. The average number of reads per sample was approximately 12,500 for the 16S rRNA gene and 20,000 (Mothur) or 34,000 (DADA2) for the 18S rRNA gene. Samples with less than 1,000 total reads were excluded from downstream analyses. Although most analyses were carried out using relative abundance, each sample was subsampled at random to normalise the number of reads per sample, and the resulting average coverage was 92% (Mothur) or 94% (DADA2) for the 16S rRNA gene and 99% (Mothur and DADA2) for the 18S rRNA gene.

### Microbial isolation and characterisation

Microbes were isolated from the final generation of positive selection experiments by plating serial dilutions on Marine Broth 2216 (BD Difco™) and mineral medium plates (*i.e*. custom medium; Supplementary information) supplemented with 0.1% N-acetyl-D-glucosamine (GlcNAc) and 1.5% agar. Colonies were re-streaked on fresh agar plates until pure isolates were obtained. The identification of isolates was carried out by sequencing the partial 16S rRNA gene (GATC BioTech, Germany) using primers 27F and 1492R (70) (Supplementary information).

Isolates were grown in custom mineral medium supplemented with either 0.1% chitin or 0.1% GlcNAc (w/v), as sources of carbon and nitrogen, to test for chitinase activity and chitin assimilation, respectively. Growth was monitored over 14 days by measuring: i) chitinase activity (as described above), ii) optical density at 600 nm, and iii) protein content (following manufacturer’s instructions; QuantiPro™ BCA Assay Kit, Sigma Aldrich, UK). Isolates were also tested on custom mineral medium agar plates made with the addition of 0.1% chitin and 0.8% agarose. Plates were incubated at 30°C for 21 days to allow the formation of halos indicative of chitinase activity.

### Statistical analyses

All analyses of chitinase activity and most MiSeq data analyses were carried out using custom Python scripts (Python versions 2.7.10 and 3.6.6) using the modules: colorsys, csv, heapq, matplotlib, numpy, os, pandas, random, scipy, scikit-bio, sklearn (71), and statsmodels. SIMPER analyses and plotting of phylogenetic trees were performed in R (R version 3.3.3) (72) using the following packages: ape (73), dplyr, ggplot2, gplots, ggtree (74), lme4, phangorn (75), plotly, tidyr, vegan (76), phyloseq (77). The top 5 ASVs identified in each SIMPER analyses were classified to their closest relative using a BLAST search of the GenBank database with a representative sequence. Hypothetical community functions were obtained using PICRUSt in QIIME1 (36, 78). Sequences used for phylogenetic trees were aligned using the SILVA Incremental Alignment (www.arb-silva.de) (79) and mid-point rooted maximum likelihood trees were constructed using QIIME1 (78). All scripts can be found at https://github.com/R- Wright-1/ChitinSelection.git. All sequences have been deposited in the NCBI Short Read Archive (SRA) database under Bioproject PRJNA499076.

Author contributions
RW and JCO designed the study. RW performed all experiments with guidance from JCO and MIG. RW wrote the first draft of the manuscript and all authors contributed substantially to revisions. The authors declare no conflicts of interest.

## Acknowledgements

Robyn Wright was supported by a Midlands Integrative Biosciences Training Partnership PhD scholarship via grant BB/M01116X/1, Joseph Christie-Oleza by National Environment Research Council Independent Research Fellowship NE/K009044/1 and Matthew Gibson by European Research Council grant 638631. MiSeq amplicon sequencing was carried out by the University of Warwick Genomics Facility. We also thank Etienne Low-Decarie, for introducing us to the concept of microbial community artificial selection, Danielle Green, for collecting seawater from Portaferry, Sally Hilton for amplicon sequencing help and advice and Vinko Zadjelovic, Gabriel Erni Cassola, Audam Chhun, and the rest of the Christie-Oleza group, for insightful discussions and advice throughout the project.

